# ShapoGraphy: a glyph-oriented visualisation approach for creating pictorial representations of bioimaging data

**DOI:** 10.1101/2021.04.07.438792

**Authors:** Muhammed Khawatmi, Yoann Steux, Sadam Zourob, Heba Sailem

## Abstract

Intuitive visualisation of quantitative microscopy data is crucial for interpreting and discovering new patterns in complex bioimage data. Existing visualisation approaches, such as bar charts, scatter plots and heat maps, do not accommodate the complexity of visual information present in microscopy data. Here we develop ShapoGraphy, a first of its kind method accompanied by a user-friendly web-based application for creating interactive quantitative pictorial representations of phenotypic data and facilitating the understanding and analysis of image datasets (www.shapography.com). ShapoGraphy enables the user to create a structure of interest as a set of shapes. Each shape can encode different variables that are mapped to the shape dimensions, colours, symbols, and stroke features. We illustrate the utility of ShapoGraphy using various image data, including high dimensional multiplexed data. Our results show that ShapoGraphy allows a better understanding of cellular phenotypes and relationships between variables. In conclusion, ShopoGraphy supports scientific discovery and communication by providing a wide range of users with a rich vocabulary to create engaging and intuitive representations of diverse data types.

## Introduction

Advances in biomedical imaging allow generating large amounts of data capturing biological systems at different scales ranging from single molecules to organs and organisms^1^. Inspection of individual images is not feasible when hundreds of images are acquired, particularly when they are composed of multiple layers, channels, or planes. Automated image analysis allows quantifying and analysing image data resulting in large multiparametric datasets^2,3^. Effective data visualisation is essential for understanding analysis results and unleashing the hidden patterns locked in these images^4,5^. However, visualising complex imaging data has been limited to general-purpose tools that do not take into account the structural nature of image data. Therefore, new visualisation techniques for representing multiparametric image data are desperately needed to aid data analysis and result interpretation from image data.

Due to their scalability to a large number of data points, heat maps and dimensionality reduction are the most widely used approaches for visualising high dimensional data, including image-based measurements. UMAPs and t-SNE are dimensionality reduction methods that project high dimensional data into lower dimensions to capture the difference between data points based on their placement in the dimensional space^6^. Heat maps represent quantitative information using colour hues, and when combined with clustering, provide a powerful tool for identifying patterns in the data^7^. There are not many visualisation techniques developed specifically for imaging data. Moreover, these approaches do not allow for intuitive representation that enables numerical information to be related to the biological entities being evaluated.

Glyph-based visualisation is another approach of visual design where quantitative information are mapped to illustrative graphics referred to as glyphs. They provide a flexible way for representing multidimensional data^9,10^. For example, we have previously developed PhenoPlot, a glyph-based visualisation approach for depicting cell shape data^11,12^. PhenoPlot was built as a MatLab toolbox that incorporates two ellipsoid glyphs to represent the cell and nucleus. It uses a variety of visual elements such as stroke, colour and symbols to encode up to 21 variables. PhenoPlot’s key benefit is that it allows for natural data mapping by selecting graphic features that resemble data attributes. For instance, the extent that a jagged border around the cell ellipse can be used to represent the irregularity of cell shape, and the proportion of ‘x’ symbols filling the cell ellipse can be mapped to endosome abundance^13^. The natural connection between the depicted quantitative representation and the measured phenomena makes it easier for the reader to interpret the visual information^10^. However, PhenoPlot is limited in terms of cell shape configuration as it is fixed to one ellipse and one sub ellipse. Furthermore, feature mapping was hard-coded, restricting its application to diverse microscopy data. Developing tools for creating new glyph-based visual encoding schemes is necessary for accommodate the diversity of biological data.

To support knowledge discovery tasks from microscopy data, we introduce a new paradigm termed glyph-oriented data visualisation. In this paradigm, data can be visualised by combining various glyph shapes to construct new visual encodings and graphical structures. We developed ShapoGraphy, a user-friendly web interface for creating such visualisations. To our knowledge, ShapoGraphy is the first method that allows modular glyph-based visualisation by integrating different shaped objects and custom mapping of shape properties, such as colour, symbols, stroke, and dimensions, to data attributes. The user can choose from a basic set of glyphs/shapes or draw their own. The effectiveness and utility of ShapoGraphy is illustrated by using various image datasets, to demonstrate how it facilitates the understanding of cellular phenotypes and interactive exploration of the data. In summary, ShapoGraphy allows the users to construct an infinite number of glyph-based representations in order to generate a quantitative and intuitive visualisation of multiparametric data supporting knowledge discovery from bioimage data.

## Results

ShapoGraphy is based on creating a template composed of one or more shapes. The various properties of each shape can be dynamically mapped to numerical data. The user has the option of selecting from a collection of predefined geometrical shapes or drawing their own (Fig. 1). Then they can position these objects relative to each other to create the desired structure (Fig. 1B). Some features can be global (the same for all data points) or dynamic (mapped to a variable). For example, the object dimensions can be changed in the global features menu or determined based on the selected variable values (Fig. 2). Other global features that can be specified include angle (rotation), fill colour, stroke colour, and opacity. For example, the user can create a cell template by adding an object for the cell body and another for the nucleus.

**Figure 1.**
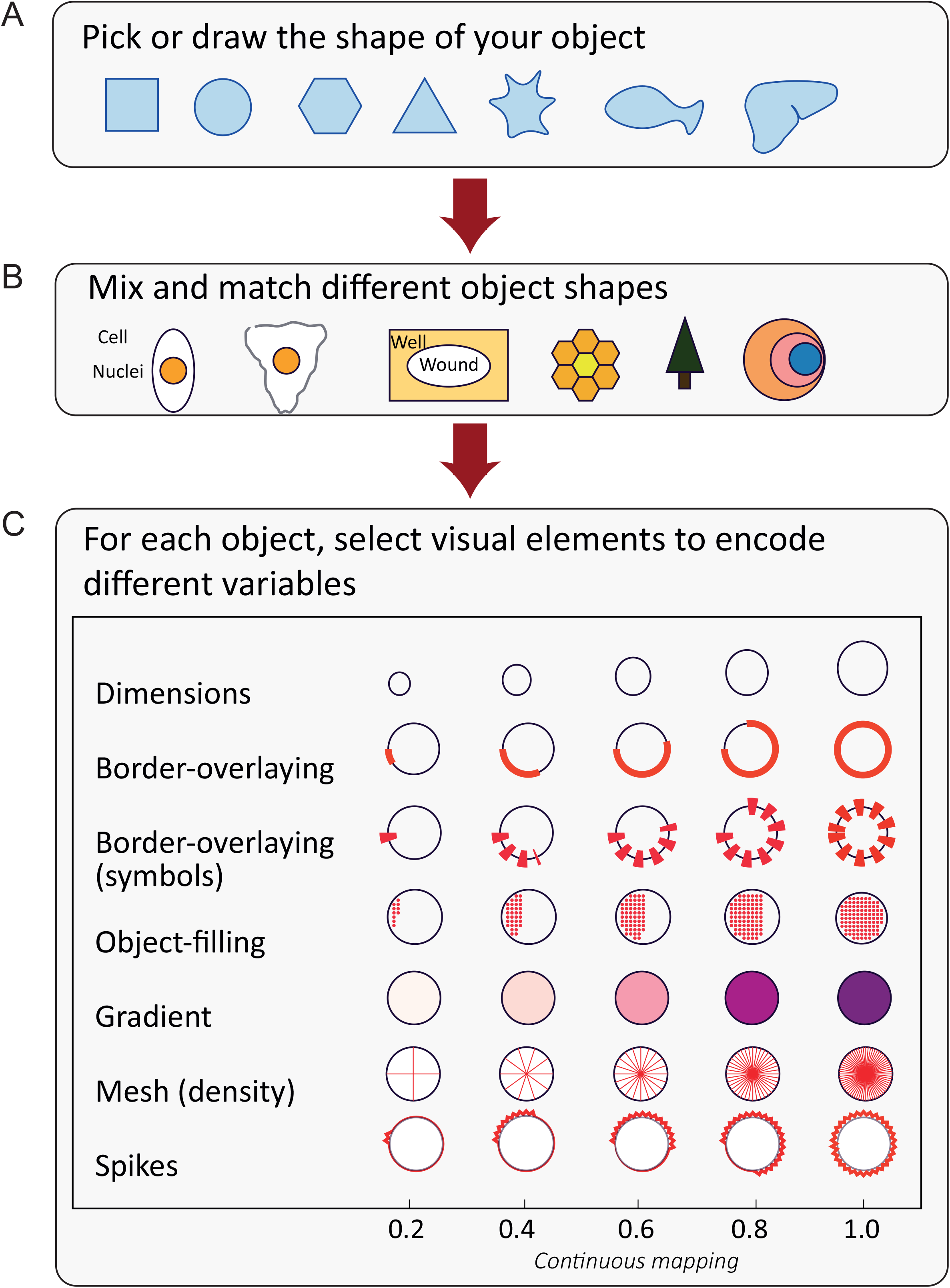
ShapoGraphy provides a highly flexible workflow for creating glyph-based visualisations.

**Figure 2.**
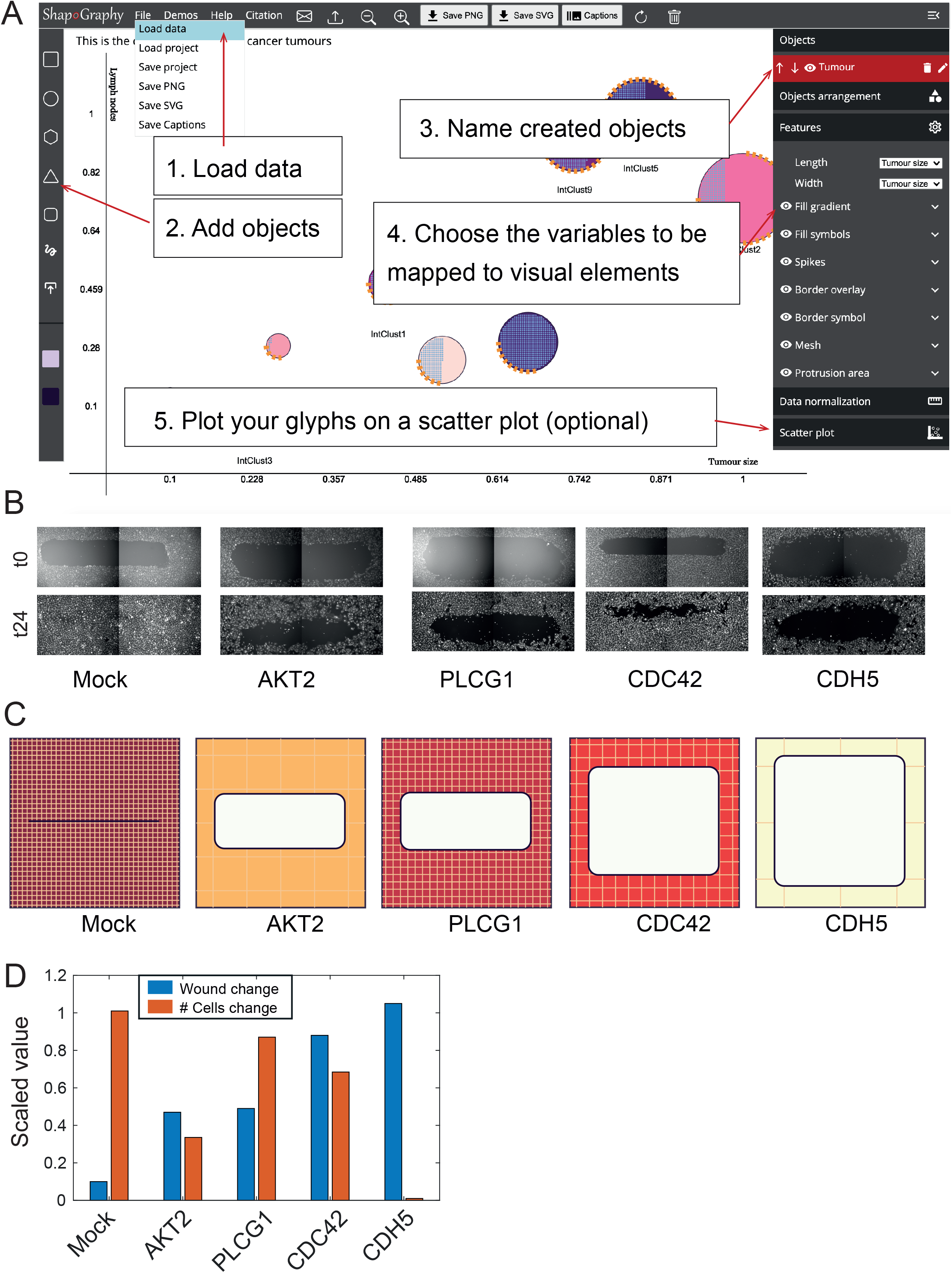
ShapoGraphy web-application. A) ShapoGraphy allows users to interactively construct and customise their plots using a flexible graphical user interface. The user creates objects and customises their properties by mapping them to the variables in their datasets. B) Image data capturing the effect of various gene depletions on human dermal lymphatic endothelial cells ability to migrate in scratch wounds (timepoint 0h and 24h). C) Intuitive representation of wound area and cell number measurements using ShapoGraphy based on data in (B). The outer square represents the well where lighter red hues indicate lower changes in cell number while higher red hues indicate higher changes in cell numbers. Change in cell number is also represented as grid density for comparison. The height of the inner square represents the normalised wound area. D) Representation of the same data in (C) using bar charts where numerical data are mapped to the bars’s length.

In order to encode continuous quantitative data to shapes, various encodings that make use of different visual elements were developed (Fig 1C). These include dimensions, size, and colour that are commonly used in graphical applications. We have previously proposed novel visual elements, such as border overlaying or object filling with symbols proportional to the variable value^11^. We introduce new features in ShapoGraphy, such as the mesh density (horizontal, vertical or grid), opacity, and rotation angle (Fig. 1C). When the users load their data, it will be scaled between 0 and 1, as in heat maps and many other glyph-based visualisations^11^. The use of various glyph shapes, positions and visual elements allows designing abstract and intuitive representations of a broad range of structures measured in biomedical imaging to assist understanding, summarising and communicating analysis results. This type of design gives the user high flexibility when it comes to constructing new visual encodings that are more intuitive and engaging.

As a first use case, ShapoGraphy was used to represent data from wound scratch assays to visualise the effects of different gene knockdowns on human dermal lymphatic endothelial cells migration into scratch wounds^14^ (Fig. 2B-D). The selected genes were shown to affect cell migration^15^. We also visualised cell proliferation because proliferation rate can also affect wound closure (Methods). We visualised the effects of different gene knockdowns using siRNA on cell proliferation and wound closure using a bar chart (Fig. 2D). The length of the bars does not allow for effective visualisation of the relationship between these two variables. ShapoGraphy, on the other hand, can be used to visualise these two variables more intuitively by representing the well and the wound as squares, where the colour of the well represents the cell number and the height of the second square represents the normalised wound area. Such representation reveals more readily that although depletion of AKT and PLCG1 genes result in a similar wound area, AKT siRNA decreases cell number. In contrast, PLCG1 siRNA increases cell number. Studying these two variables can lead to a different interpretation of their effect on cell migration. Similarly, depleting CDH5 and CDC42 significantly affect cell migration, but CDC42 siRNA increases cell number while CDH5 siRNA reduces it (Fig. 2C). Such information are difficult to discern from raw images as wound measurements need to be normalised to timepoint 0h (Fig. 2B). These results show how ShapoGraphy can be used to understand biological phenotypes better and identify relationships between variables.

Next, ShapoGraphy was used to illustrate how it can assist the interpretation of single cell phenotypes in high dimensional multiplexed imaging data measuring 40 markers^16^. Multiplexed imaging allows simultaneous capture of spatial protein activities, subcellular organisation as well as various cell identities^17^. Since tens of markers can be imaged, colour coding of the different proteins is no longer useful to visualise this information^1^. To study the phenotypic heterogeneity of cancer HeLa cells, we analysed data from 2000 cells that were stained with markers highlighting various cellular organelles and signalling components including mTOR pathway (Fig. 1A and Methods). Cells were clustered and visualised using UMAP to characterise different subpopulations in the data (Fig. 3B and Methods). Although this analysis provides a high-level picture of the data structure, the underlying phenotypic differences between clusters cannot be obtained. Heat maps allow studying all the measured markers individually but require visual search by the user, memorisation and many cognitive calculations to build a picture of what these clusters represent (Fig. 3A). Since the coloured rectangles do not incorporate the semantics of the data, they can be challenging to interpret and require memorisation of multi-protein values.

**Figure. 3.**
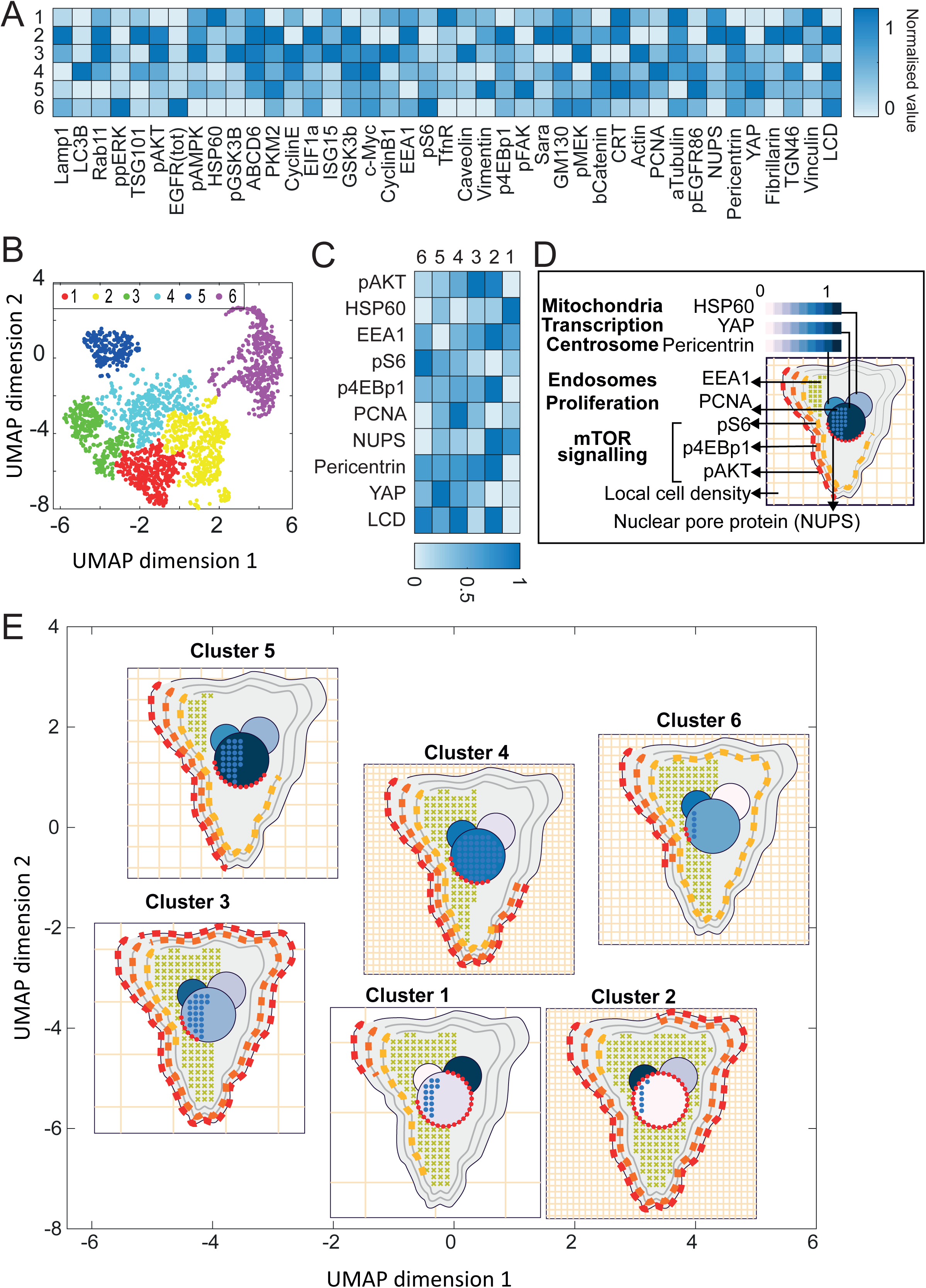
ShapoGraphy allows interpreting multiplexed single-cell data. A) Representation of average values of 40 markers values as well as local cell density for each identified cluster using a heat map. B) UMAP projection of 2000 single HeLa cells. C) Selected features from A based on qualitative exploration. D) Representation of the data in C using ShapoGraphy where the Shape Graphs are placed based on cluster centre. Shape Graphs allow a better understanding of differences and similarities between clusters, and better memorisation by providing natural mapping of the plotted data attributes.

In order to facilitate the understanding of single cell phenotypes that are derived from multiplexed data, ShapoGraphy was used to design a template where the visual elements resemble the represented data attributes (Methods). Furthermore, we positioned the composite glyphs based on the centre of identified clusters in the reduced UMAP space to help sorting these composite glyphs and comparing cluster phenotypes (Methods). Figure 3D shows that Cluster 2, 4, and 6 on the right have high cell density (grid density). Cluster 6 and 4 are highly similar, but Cluster 6 has the highest pS6 levels across all clusters, while Cluster 2 has very high pAKT and p4EBp1, centrosomes abundance (Pericentrin), nuclear pore proteins (NUPs), early endosomes (EEA1) and low YAP values. Cluster 3 also has high pAKT and p4EBp1 like Cluster 2 but has lower density and higher YAP values. Furthermore, we observe that pAKT and p4EBp1 levels are correlated across clusters but not pS6. Discussing our results with biologists, they found that these representations help them understand their data better. In comparison, Figure 3C depicts the same information in a heat map which can complement our shape graphs but does not help the reader to build a mental picture of the data. Therefore, ShapoGraphy allows a more intuitive representation of phenotypic classes and their biological relevance based on high dimensional single-cell data.

## Discussion

The human brain perceives information by converting visual stimuli to symbolic representations that are then interpreted based on our memories and previous knowledge. Visualisation approaches help our brain to create a mental visual image of quantitative data in order to recognise patterns and identify interesting relationships that might otherwise go unnoticed^18^. ShapoGraphy is a new visualisation approach that allows the user to create representations that mimic the measured phenomena by constructing a custom collection of shapes that can encode multiple variables. To our knowledge, such an approach to data visualisation has not been explicitly proposed before and no tool is available to create such graphical representations automatically. The main advantage of ShapoGraphy is that it enables generating a metaphoric realistic quantitative representation of the data to aid the reader in creating a mental visual image that improves interpretation, understanding, memorisation and communication of the data. Another advantage of Shape Graphs is that such pictorial representations can attract more attention by the reader as they stimulate more cognitive activities^9^. This is very important when the goal is to communicate the data with a wider range of audience. Therefore, ShapoGraphy serves as a general-purpose methodology for creating more engaging and intuitive graphic templates.

ShapoGraphy complements existing visualisation methods such as heat maps, t-SNE and UMAPs. While the later approaches provide a global picture of the major trends or structure in the data, ShapoGraphy allows more detailed understanding of multiparametric phenotypes. Notably, ShapoGrapy aims to represent quantitative data so the user can compare different variable values relative to each other, rather than generating an actual abstract picture of the image data. Such distinction is necessary as image data are often normalised which make interpreting raw image data more challenging and subjective. Currently our approach is best suited for summarising major phenotypes in the data because of the pictorial nature of the generated representations, which require high resolution. These phenotypes can be identified using clustering or classification tasks. A potential future direction is to extend our approach to gain multi-level summaries of the data enabling visualisation of a larger number of data points.

One important aspect when composing different shapes is the consideration of Gestalt perceptual principles that state that we tend to group objects and perceive visual components as a whole or as organised patterns^19,20^. These principles include proximity, similarity, continuity, closure, figure/ground, area, and symmetry. For example, humans tend to group objects closer to each other or auto-complete an incomplete shape. In accordance with the figure/ground principle, less important objects can be assigned a lighter colour, while more important objects can be assigned darker colours. It is also desirable to minimise occlusions when choosing various visual elements, and use the same colour scheme for adjacent objects to enable comparison. Another important factor in designing effective visualisation is to remove irrelevant information as our working mental memory is limited and can handle only a few variables (5-10) at a time^4^. Selection of important features can be achieved through interactive exploration to identify the most relevant information to be communicated with the reader. Notably, new visual encodings to represent specific or complex domain knowledge can require time to learn^10^. Once learned, such glyph-based visualisations have been shown to be useful in all fields of biology as they provide richer context.

To conclude, ShapoGraphy can be used in all steps of data analysis to create intuitive pictorial representations of any data type. It can be used to summarise analysis results obtained from clustering or classification approaches, as well as an educational tool. We believe that the unique flexibility offered by ShapoGraphy will expand our visual vocabularies, accelerate the evolution of glyph-based visualisation and stimulate the development of new visual encoding schemas.

## Methods

### Software

ShapoGraphy is a client-side web application developed using HTML5 and JavaScript (www.shapography.com). This means that all the processing happened at the user end and minimal data is uploaded to our server. This makes our tools highly efficient and circumvents privacy issues.

We defined a portfolio of templates to accommodate different data types. The user can choose an existing template to map their data. The user can modify an existing template or delete unwanted objects for a maximised flexibility. The created graphical templates and data mapping by ShapoGraphy can be saved to be used later.

### Datasets

#### Wound scratch data

Wound scratch data was obtained from an image-based siRNA screen measuring human dermal lymphatic endothelial cells migration into a scratch wound created in a cell monolayer^20^. Cells were imaged at 0h and 24h following wounding at 4x objective. Cells were detected and the wound area was segmented using DeepScratch^15^. Measurements of wound size and cell numbers at 24h were normalised to timepoint 0h and represented using ShapoGraphy.

#### Multiplexed imaging data

Multiplexed imaging data of 2000 HeLa cells was obtained from Gut et al., 2018 where immunofluorescence of different markers was performed in cycles to image the subcellular localisation of 40 proteins^16^ (Fig. 3A). Data was scaled and the number of dimensions were reduced using UMAP. K-means was used to group phenotypically similar cells into six clusters.

The average of each cluster was represented using ShapoGraphy. Three cell-shaped objects were created to represent PI3K/AKT/mTOR pathway (pAKT, p4EBp1 and pS6) on the cell periphery as the proportion of symbols overlayed on the object border (Fig. 3D). The grid density in the square surrounding the cell represents the local cell density. The abundance of early endosomes (EEA1) was represented as ‘x’ symbols filling the cytosol. Mitochondria and centrosome organelles were abstracted as circles with a colour gradient reflecting their abundance. Three variables were mapped to the circle-shaped nucleus: the value of nuclear pore protein (NUPs) was mapped to the nuclear membrane, the level of YAP transcription factor was mapped to the colour of the circle, and the abundance of cell proliferation protein PCNA was represented as circle symbols filling the nucleus. Using this representation, it is easier to connect the different variables to the relevant biological entities and build a mental image of the data. The position of each data point is mapped to the cluster centre using the positional mapping sub-menu.

## Acknowledgments

HS is funded by a Sir Henry Wellcome Fellowship (Grant Number 204724/Z/16/Z).

